# NCOR1+/CD4+ T cells in tertiary lymphoid organs suppresses tumor growth and predicts favourable prognosis in pancreatic ductal adenocarcinoma

**DOI:** 10.1101/2023.10.15.562444

**Authors:** Tiangeng Dong, Tuo Yi, Yuda Gong, Weidong Gao, Bo Zhang, Weizhong Sheng

**Affiliations:** Department of General Surgery, Zhongshan Hospital, Fudan University, Shanghai, China

**Keywords:** pancreatic cancer, NCOR1, CD4 T cells, tertiary lymphoid organs

## Abstract

CD4+ T cells have critical roles in anti-tumour immunity and its differentiation is known to be regulated by the nuclear receptor co-repressor 1 (NCOR1). Previous studies suggested that high CD4+ T cells are associated with a favourable prognosis in patients with pancreatic ductal adenocarcinoma (PDAC). However, the prognostic significance of NCOR1 in PDAC is still missing. In this study, the pathological impact of NCOR1 and CD4 has been analysed by multiplex immunohistochemistry in 100 PADC patients. NCOR1 expression in tertiary lymphoid organs is positively associated with the good prognosis of PDAC patients, while its expression in tumour tissue is not. Furthermore, the presence of NCOR1+/CD4+ T cells in tertiary lymphoid organs predicts a favourable prognosis in PDAC. Mechanistically, upregulation of NCOR1 expression in CD4+ T cells increases the release of TNF-α, which induces the apoptosis of the tumor cells *in vitro*. Together, our data highlighted the tumour suppressive role of NCOR1+/CD4+ T cells in PDAC.

## Introduction

Pancreatic ductal adenocarcinoma (PDAC) is a common gastrointestinal malignancy with a poor prognosis (1). The overall 5-year survival rate of PDAC is 3–5%, and that of resectable patients is only 15-25% (1). Conventional chemical-based therapies for PDAC are ineffective to prolong the patients’ survival with frequent occurrence of chemoresistance and often intensify the physical and mental suffering of patients (2, 3). Novel treatments such as targeted therapies bring great hope to PDAC patients (2). Discovering biomarkers to assist the selection of patient subsets is useful for clinical purposes, especially for evaluating therapeutic approaches and developing novel treatments.

The tumour microenvironment has been recently recognized as an important factor in tumorigenesis (4). Among the tumour microenvironment, the tumour infiltrating lymphocytes (TILs) has been found to have prognostic value in PDAC patients and other solid tumours such as colorectal, lung and breast cancers (5, 6). TILs consist of a heterogeneous population of lymphocytes e.g. dendritic cells, macrophages, CD4+ T cells and CD8+T cells etc. TILs are known to promote tumour progression or battle against tumorigenesis. For example, Th1 CD4+ T cells, cytotoxic CD8+ T cells, and natural killer cells tend to inhibit tumour growth, while Th2 and Th17 CD4+ T cells, and FOXP3+CD4+ regulatory T cells (Tregs) are regarded as tumour-driving factors dependent on tumour types. In PDAC patients, high numbers of CD8+ TILs and CD4+ TILs had a significantly longer overall survival (OS) rate than patients with low numbers (7). In addition, the high density of FOXP3+ Tregs is correlated with lymph node metastasis and poor prognosis in PDAC patients (8).

Differentiation of CD4+ T cells is tightly controlled since dysregulation can result in autoimmune-associated diseases such as allergies (9). Cytokine signalling along with activation of lineage-specific transcription factors and epigenetic modifications are required for its differentiation (10, 11). Among the cofactors that are involved in differentiation, nuclear receptor corepressor (NCOR1) is recently found to facilitate the formation of repressive chromatin-modifying enzymes e.g. histone deacetylases (HDACs) and transcription factors (12). NCOR1 is a non-DNA-binding protein that mediates transcriptional repression of nuclear receptor and also interacts with many other types of transcription factors (13). NCOR1 plays an important role in muscle physiology, liver metabolism and immune regulation (14-16). The role of NCOR1 in T cell differentiation has been recently studied. Several studies revealed that NCOR1 was able to promote the survival of activated T cells (17, 18). In particular, NCOR1 deletion mostly affect the CD4+ T cell activation and Th1/Th17 effector functions (19). These studies suggested that NOCR1 is an important regulator for the transcriptional activation in naive and effector CD4+ T cells. Here, we further determined the clinical significance of NCOR1 and CD4+ T cells in predicting the survival of PDAC patients and also the function of NCOR1 in inhibiting PDAC growth.

## Materials and methods

### Patients and sample collection

We enrolled 100 patients who underwent R0 resection for PDAC identified by the pathology department at the Zhongshan Hospital (Shanghai, China). All the patients enrolled had resectable PDAC, which had no arterial tumour contact (celiac axis [CA], superior mesenteric artery [SMA], or common hepatic artery [CHA]); and no tumour contact with the superior mesenteric vein (SMV) or portal vein (PV) or less than 180° contact without vein contour irregularity, and without distant metastasis. Overall survival (OS) was calculated as the interval between the date of surgery and the date of death or the last follow-up visit. Recurrence-free survival (RFS) was defined as the interval between the date of surgery and the date of tumour recurrence or the last follow-up visit. All patients were followed up until December 2021. The use of human PDAC tissues was approved by the Research Ethics Committees at Zhongshan Hospital, Fudan University. Informed consent was obtained from all patients according to the committees’ regulations.

### Multiplexed immunohistochemistry

Multiplex immunofluorescence staining was performed using PANO 4-plex IHC kit! Cat 10004100100 (Panovue, Beijing, China). CD4, Foxp3 and NCOR1 antibodies were sequentially applied, followed by horseradish peroxidase-conjugated secondary antibody incubation and tyramide signal amplification. The slides were microwave heat-treated after each TSA operation. Images of unstained and single-stained sections were used to extract the spectrum of autofluorescence of tissues and each fluorescein, respectively. Nuclei were stained with DAPI after all the human antigens had been labelled. To obtain multispectral images, the stained slides were scanned using the Mantra System (PerkinElmer, Waltham, Massachusetts, US), which captures the fluorescent spectra at 20-nm wavelength intervals from 420 to 720 nm with identical exposure time; the scans were combined to build a single stack image. The extracted images were further used to establish a spectral library required for multispectral unmixing by inForm image analysis software (PerkinElmer, Waltham, Massachusetts, US).

### Lentivirus packaging and transfection

For constructing NCOR1 overexpression CD4 T cells, we used the lentivirus transaction strategy. The virus was packaged by the lipofection of three vectors together, the packing vector PAX2, the envelop vector pMD2G, and the transfer vector pCMV carrying the ORF of NCOR1 gene. Lipo3000 (Thermo Fisher) was used for packing as described by the manufacture’s protocol. Lentivirus was harvested at 24h, 48h post lipofection, and then used to transfect the CD4 T cells. For isolation of CD4 T cells, we used the EasySep™ Human CD4+ T Cell Isolation Kit (Stemcell). Briefly, fresh blood was acquired from an PDAC patient and separated by centrifugation to acquired the PBMCs. The following steps were conducted as described by the manufacture’s protocol.

### Cell culture

The human PDAC cell-line Panc1 was cultured in DMEM supplemented by 10% fetal bovine serum. CD4 T cells were cultured in RPMI 1640 basal medium supplemented with 10% fetal bovine serum. CD4 T cells was added directly to the Panc1 for co-culture assay.

### Tunnel assay

For detecting the apoptosis of Panc1 cells, TUNEL Assay Kit(Cell signling #48513) was used. The culture medium was firstly removed and the Panc1 was washed by 1×PBS twice to remove the CD4 T cells. Then, Panc1 was trypsinized to achieve the disperse single cells. The following steps were followed as described by the manufacture’s protocol. Flow cytometry was used to analyze the positive cells by using the excitation 593 nm / emission = 614 nm.

### RT-qPCR analysis

Total RNA was isolated by Trizol. Quantification was performed by The LightCycler 480 (Roche). RNU6-1 was used as the control. Reverse transcription was performed by using FastQuant RT kit. cDNA samples were prepared in Hieff qPCR SYBR Master Mix (YEASEN) and quantified by The LightCycler 480 (Roche).

### Western-blot analysis

Samples were collected in 1 × protein loading buffer, and total proteins were separated by SDS polyacrylamide gel electrophoresis and transferred onto the NC membrane. The membrane was blocked using 1 × blocking buffer and incubated with the indicated primary antibody and secondary antibody. Bands were visualized using GE AI600.

### Statistical analysis

OS and RFS were displayed on Kaplan–Meier survival curves with 95% confidence intervals (CIs). P < 0.05 was considered statistically significant. Statistical analyses were performed using SPSS 24.0 (SPSS Inc., Chicago, IL).

### Results and discussion

Firstly, we performed an immunohistochemistry analysis of the NCOR1 expression in 100 PDAC samples from resectable patients. Although NCOR1 expression could be detected in tumour tissues with different levels (Figure 1A), its expression in tumour tissues showed no significant correlation with the patients’ overall survival and relapse-free survival (Figure 1B, Figure 1D). Besides NCOR1 expression in tumour tissues, an obvious expression could be seen in tumour-neighbored tertiary lymphoid organs (TLOs) (Figure 1A). TLOs “also called lymphoid organ-like structures or ectopic lymphoid organs# are found at the sites of chronic inflammation in autoimmune diseases and are even often identified in human tumours (20, 21). Recent studies suggested that TLOs might be an important modulator of the cancer immunological microenvironment. Therefore, we analysed the correlation between NCOR1 expression in TLOs and patients’ survival. Strikingly, a high level of NCOR1 in TLOs was positively associated with a good prognosis (Figure 1C, Figure 1E). Of note, more TLOs suggested a better prognosis in PDAC (data not shown), which is consistent with a previous study that the intratumoral TLO is a favourable prognostic marker in PDAC (22).

**Figure1.**
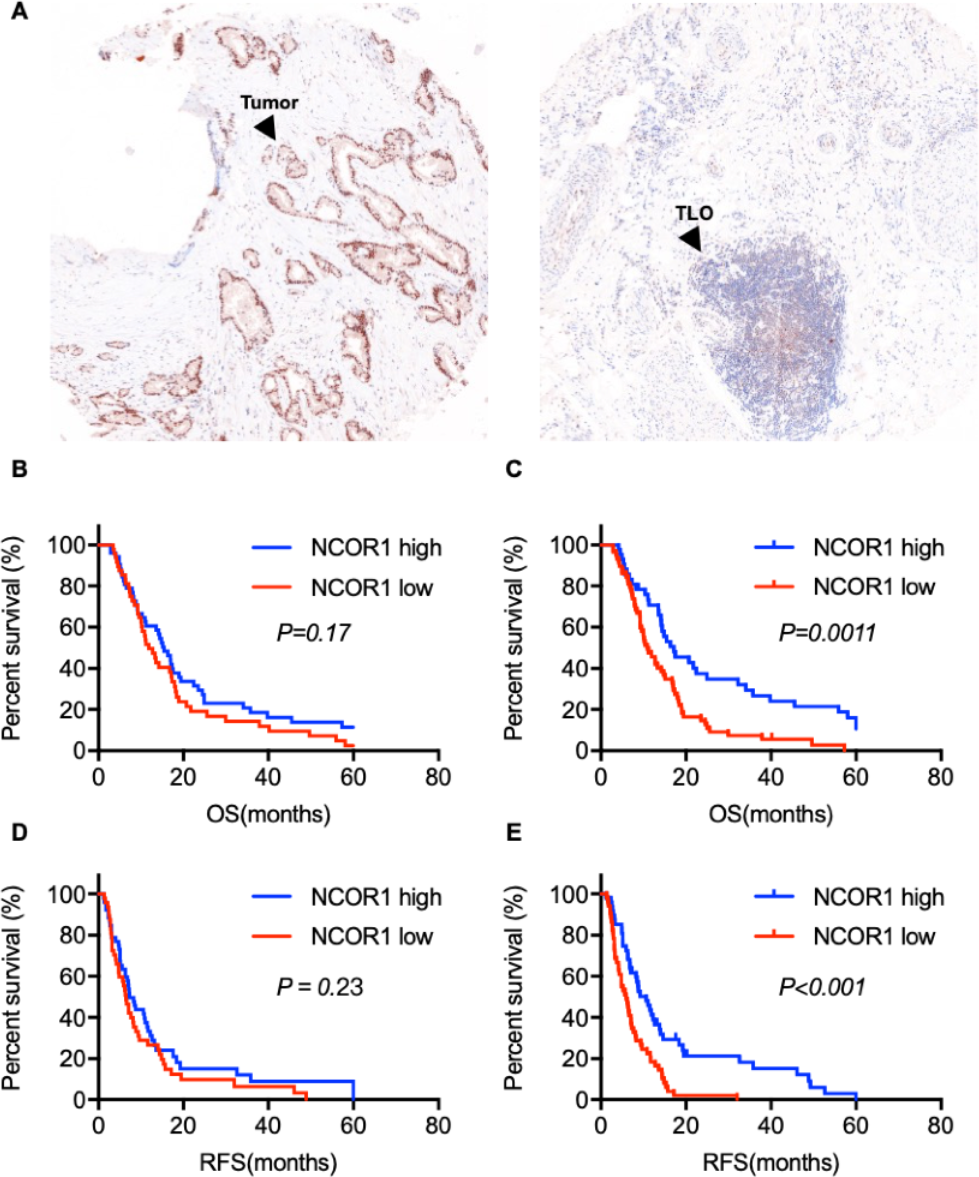
NCOR1 expression in tertiary lymphoid organs is a prognostic marker in PDAC. (A) A represented image of NCOR1 expression in PDAC tissues analyzed by immunohistochemistry. (B) and (C) Statistics of PDAC patients’ overall survival (OS). (D) and (E) Statistics of PDAC patients’ relapse-free survival (RFS).

Given that NCOR1 has a critical role in CD4+ T cell differentiation and there is an abundant distribution of CD4+ T cells in TLOs, we performed multiplex immunohistochemistry to simultaneously analyze the expression of CD4, NCOR1 (Figure2). Notably, we also detected the Foxp3 expression, which is a maker of regulatory T cells (Figure2). We found that the regulatory T cells (CD4+/Foxp3+) level in PDAC tumours was negatively associated with the good prognosis of PDAC patients (data not shown), which was in agreement with previous studies (23, 24). The distribution of CD4+ T cells could be observed in the PDAC tumours, the neighbored tissues, as well as in the TLOs. As expected, NCOR1+/CD4+ T cells presented a high abundant distribution in TLOs in patients with better survival. Significantly, statistical analysis demonstrated that a high percentage of NCOR1+/CD4+ T cells predicted a good prognosis in PDAC patients (Figure 2B, 2C). Together, our results suggested that infiltration of NCOR1+/CD4+ T cells in TLOs in PDAC tumour tissues may facilitate the anti-tumour effects.

**Figure2.**
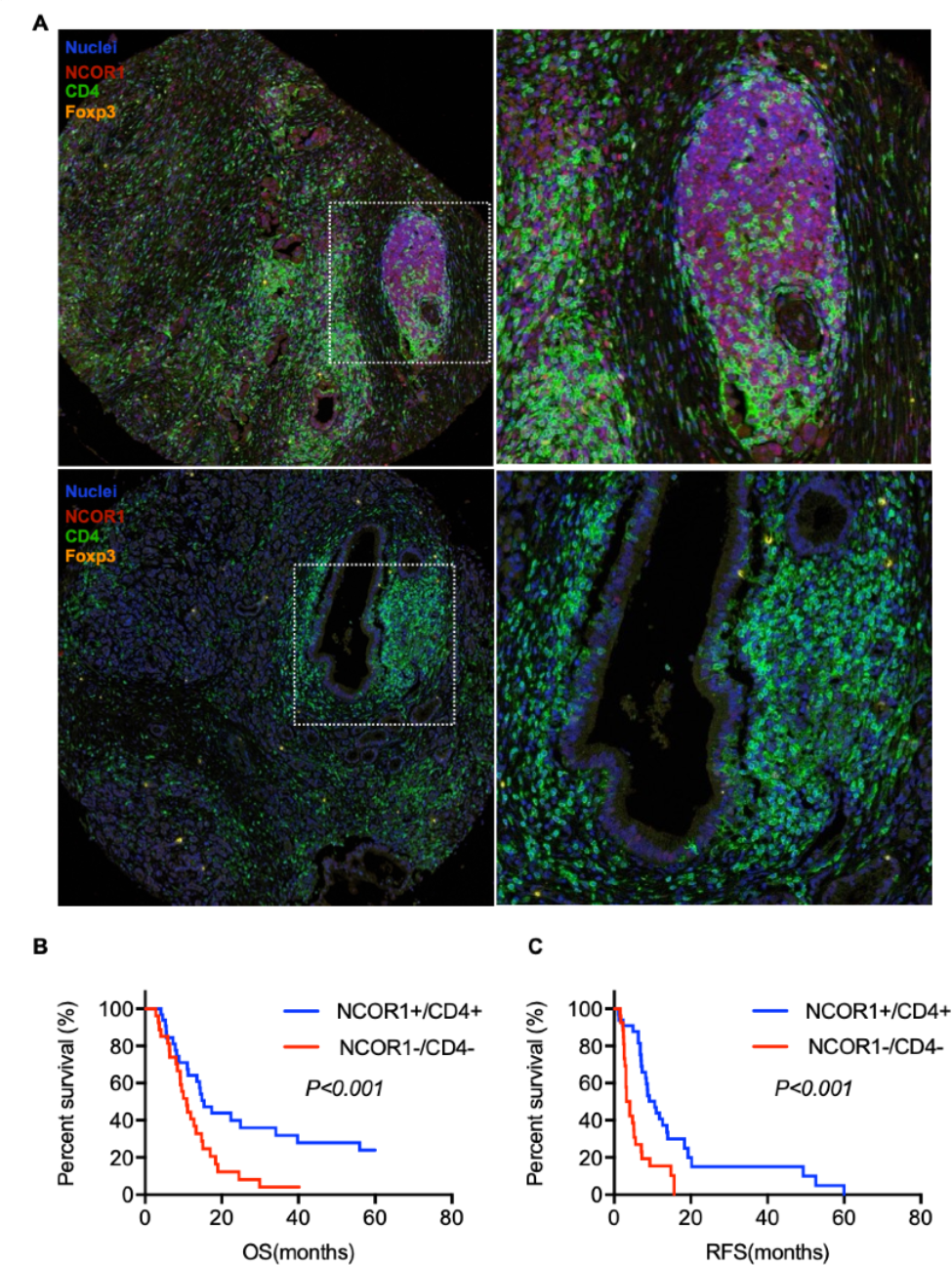
NCOR1+/CD4+ T cells in tertiary lymphoid organs is a well prognostic marker in PDAC. (A) A represented image of NCOR1, CD4, Foxp3 expression in PDAC tissues analyzed by multiple-immunohistochemistry. (B) Statistics of PDAC patients’ overall survival (OS). (C) Statistics of PDAC patients’ relapse-free survival (RFS).

It is well-known that CD4+ T cells play an important role in anti-tumour immunity by assisting the priming of tumour-specific cytotoxic T lymphocytes in lymphoid organs (25). In PDAC, CD4+ T cells were recognized as a prognostic marker in several studies (26, 27). However, there are versatile subtypes of CD4+ T cells and a more accurate deification of CD4+ T cells for predicting PDAC prognosis is still lacking. Here, we further determined that the abundance of NCOR1+/CD4+ T cells in TLOs in PDAC tumours was a favourable prognostic marker. Our data suggested that the NCOR1 may play an unknown role in the CD4+ T cell differentiation in TLOs.

Recent studies suggested that CD4 T cells affect tumorigenesis by direct or indirect mechanisms (28). Next, we decided to study the biological function of NCOR1 in CD4 T cells in tumor progression. Direct mechanisms of CD4 T cell-mediated anti-tumor activity included the cytokine release of CD4 T cells activated by tumor cells, especially TNF-α and IFN-γ (28). We isolated the CD4 T cells from PBMCs (Peripheral Blood Mononuclear Cells) of an PDAC patient and constructed NCOR1 overexpression CD4 T cells by lentivirus transfection as described in the methods. The NCOR1 overexpression CD4 T cells were successfully acquired and determined by western-blot analysis (Figure 3A). TNF-α level were significantly unregulated at the transcriptional level while no difference could be observed about the IFN-γ level (Figure 3B), suggesting that NCOR1 majorly affect the TNF-α pathway. In consistent with the RT-qPCR data, the released TNF-α was unregulated in NCOR1 overexpression CD4 cells, while no alteration of released IFN-γ were detected by Elisa assay (data not included). Co-culture of CD4 T cell and Panc1 cells were conducted as described in methods (Figure 3C). Compared to the control, where empty indicated no additional element for a negative control and TNF-α (Invivo Gen, working concentration was 0.1 ug/mL) was added for a positive control, co-culture of NCOR1 overexpression CD4 T cells reduced the tumor cell proliferation to approximately 30% (Figure 3D). Corresponding to this result, the apoptosis of Panc1 was significantly activated by the co-culture of NCOR1 overexpression CD4 T cells (Figure 3E). Of note, we used mono-clonal anti-TNF-α to block the effect of TNF-α, which only reverse the tumor cell proliferation to about 70% (data not shown), suggesting that NCOR1 may also regulate other factors to affect tumor proliferation. Future investigation is needed to clarify the complicated downstream pathway following NCOR1 overexpression in CD4 T cells.

**Figure3.**
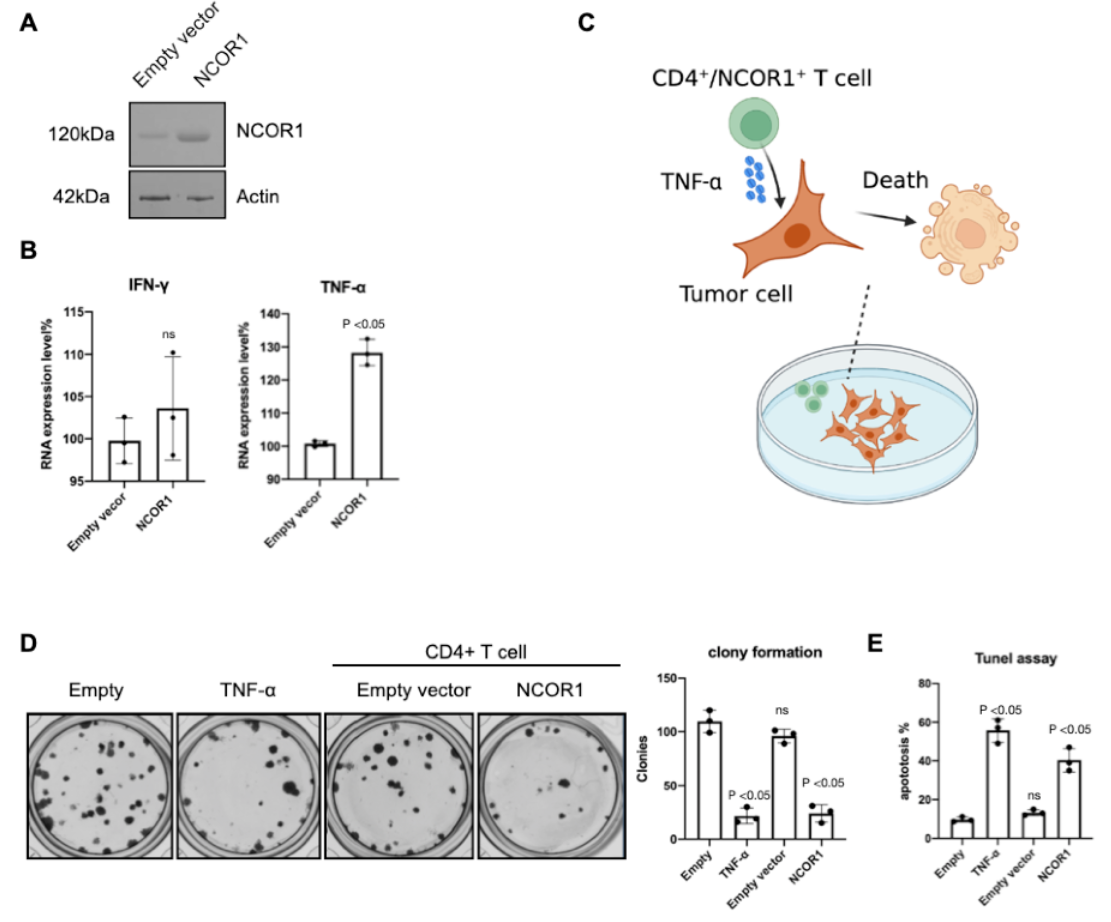
NCOR1 overexpression in CD4+ T cells induce the release of TNF-α and thus inhibit tumor cell proliferation. (A) Western-blot analysis CD4 T cells transfected with indicated lentivirus. (B) RT-qPCR analysis of gene expression. Data were normalized by comparison to the GAPDH level. (C) Schematic representation of assay setup for the co-culture of CD4 T cells and PDAC cell-line Panc1. (D) Colony formation assay of Panc1. (E) Tunnel assay for apoptosis quantification. Results were calculated by the percentage of total events. All data were performed at least three independent assays.

Together, our results demonstrated that NCOR1+/CD4+ T cells level in PDAC was a prognostic maker and further investigation is needed to clarify the functional role of NCOR1+/CD4+ T cells in PDAC tumorigenesis and tumour immunity.

## Notes

### Competing Interest Statement

The authors have declared no competing interest.

